# Identification of Chikungunya virus nucleocapsid core assembly modulators

**DOI:** 10.1101/774943

**Authors:** Sara E. Jones-Burrage, Zhenning Tan, Lichun Li, Adam Zlotnick, Suchetana Mukhopadhyay

## Abstract

The alphavirus Chikungunya virus is transmitted to humans via infected mosquitos. Most infected humans experience symptoms which can range from short-term fatigue and fever to debilitating arthritis that can last for months or years. Some patients relapse and experience symptoms months or years after the initial bout of disease. The capsid protein of Chikungunya virus forms a shell around the viral RNA genome; this structure is called the nucleocapsid core. The core protects the genome during virus transmission and with the correct environmental trigger, this proteinaceous shell dissociates and releases the viral genome to initiate infection. We hypothesized that targeting compounds to interfere with the nucleocapsid core’s function would constrain virus spread either by inhibiting the release of viral genomes during entry or by reducing the number of infectious virus particles assembled. We implemented a high throughput, *in vitro,* FRET-based assay to monitor nucleic acid packaging by purified Chikungunya capsid protein as a proxy for nucleocapsid core assembly and disassembly. We screened 10,000 compounds and found 45 that substantially modulated the assembly of core-like particles. A subset of compounds was selected to study their effects in virus-infected vertebrate cells. Our results show that four compounds inhibit infectious virus production by at least 90% in a dose-dependent manner. The most promising inhibitor was tested and found to reduce the amount of nucleocapsid cores inside the cell during Chikungunya virus infection. These compounds could be the foundation for anti-viral therapeutics.

**Highlights:**

- A FRET-based assay to detect nucleic acid packaging by Chikungunya virus capsid protein
- Identification of small molecules that modulate core-like particle assembly
- A subset of compounds that interfere with in vitro assembly also inhibit Chikungunya virus production in cell culture
- Identification of antiviral molecules that may not be identified by assays using reporter viruses
- Potential starting compounds for developing direct-acting antivirals

## 1. Introduction

Chikungunya virus (CHIKV) is a re-emerging alphavirus that is spread by a mosquito vector to humans and other vertebrates (1, 2). An estimated 3 million people are affected annually by CHIKV and approximately 80-95% develop symptoms 2 to 12 days after being bitten by an infected mosquito (1, 3, 4). Infected individuals typically suffer from short-term fever, fatigue, and joint pain; however, approximately 25-48% develop long-term debilitating arthritis that can persist for months or years (2, 5, 6). Previously, CHIKV outbreaks were limited to tropical and subtropical Asia and Africa, but outbreaks have now been reported in Europe, North America, South America, and the Caribbean (2). This expansion is due in part to a mutation in a viral protein, which supportstransmission of the virus in both the *Aedes aegypti* and *Aedes albopictus* mosquito vectors (7). A number of inhibitors of CHIKV have been identified including compounds that target the virus lifecycle including genome replication, viral protein synthesis, and virus egress, specifically the interaction of the core with the viral spikes (8–19). These compounds, however, are effective only at high concentrations or resistance is easily acquired. Antiviral compounds that target host factors required by CHIKV reduce viral titers but these compounds are by definition affecting normal cellular function (10, 20–22).

Antivirals that target the functions of capsid proteins of approximately half a dozen other eukaryotic viruses have been successful direct acting antivirals. While the core is modulated in all the inhibitors identified, their mode of action and lifecycle stage varies. Work with polio and dengue viruses (23, 24) show the that when antivirals (V-073 for polio and ST-148 for dengue) target the capsid/core proteins, the virus is less likely to develop drug resistance because the viral capsids/cores will contain both drug-susceptible and drug-resistant capsid proteins. Because of this chimera oligomer, the drug is still effective despite mutations arising. Additional studies have shown that ST-148, is thought to make the dengue core more rigid by promoting capsid contacts, thus interfering with the packaging and disassembly of viral RNA (25, 26). One could envision that a dengue core containing even a few core proteins that have stronger contacts with adjacent subunits would disrupt the overall assembly of the core. Small molecules have been identified that target Hepatitis B virus capsid assembly (27–31); heteroaryldihydropyrimidines compounds promote assembly and lead to empty or misassembled particles that have fallen into kinetic traps (27, 30). In contrast, inhibitors of enteroviruses, such as the WIN compounds for rhinoviruses, prevent the disassembly of the capsid particle (32, 33). Here, the two most efficient WIN compounds bind to a pocket in the capsid protein inhibiting breathing and disassembly of the capsid (33, 34). Two groups of antivirals that target HIV capsid have been identified. Small molecules will interfere with the virion maturation by targeting the cleavage and rearrangement of capsid, which is necessary for infectious viron maturation (35–42). The second group of antivirals interfere with the capsid -capsid protein interactions, and interfere with assembly, genome packaging, and uncoating (40, 43, 44).

CHIKV, like other Alphaviruses (Ross River virus, Venezuelan Equine Encephalitis virus, Eastern Equine Encephalitis virus) are enveloped, positive-sense RNA viruses (45). Particles have icosahedral symmetry and are organized with (i) an internal nucleocapsid core, consisting of the capsid protein and RNA genome, (ii) a host-derived lipid bilayer surrounding this core, and (iii) a shell of viral glycoproteins on the particle surface which are required for cell entry (46, 47). The capsid protein consists of two domains, a disordered N-terminal domain that interacts with the viral RNA, and a C-terminal chymotrypsin-like domain that makes up of the outer surface of the core. The C-terminal domain of capsid protein interacts with the endo domain of one of the viral glycoproteins, E2. The different roles of the capsid protein in the viral lifecycle require multiple interactions of capsid protein.

Assembly of alphavirus nucleocapsid cores can be recapitulated *in vitro*; these are called core-like particles (CLPs) (48). CLPs can be assembled by mixing recombinant capsid protein with viral RNA. CLPs are also formed when non-viral cargo (*e.g.,* single-stranded DNA and RNA oligonucleotides, polyanion small molecules, or gold particles coated with nucleic acid) are mixed with capsid protein (49). CLPs can be encapsidated by viral glycoproteins, and the newly formed virus-like particles are able to enter and disassemble within a new cell (50, 51). A 27mer DNA oligo interacts in a 1:1 molar ratio to neutralize the basic N-terminus of the capsid protein (52, 53) to initiate CLP assembly. CLPs are 40 nm in diameter and structurally similar to cores in virions (52–55). The efficiency of CLP assembly can be regulated by ionic strength. Low ionic strength buffers promote CLP assembly and inhibit CLP disassembly. High ionic strength buffers will inhibit CLP assembly and will promote CLP disassembly (48-50, 52-54, 56). The need for anionic cargo and the sensitivity to ionic strength indicate that the capsid protein-capsid protein interactions during assembly are extremely weak and that assembly depends on electrostatic interaction with cargo.

In this study, we sought to identify small molecule modulators of CHIKV CLP assembly (48, 53, 57). We monitored the fluorescence quenching of fluorophore-labelled oligonucleotides to monitor nucleic acid packaging. We screened a 10,000 compound library *in vitro*. From the 45 compounds identified *in vitro*, 12 were selected for validation in cell culture with CHIKV. We report that four compounds could block the production of infectious CHIKV particles by at least 90% in a dose-dependent manner. Furthermore, the compound that reduced titers the most, reduced the amount of cytoplasmic cores present in infected cells, consistent with this compound targeting the nucleocapsid core during infection.

## 2. Materials and Methods

### 2.1 Capsid and oligomer preparation

The capsid protein from CHIK 181/25 (58) was cloned into the pET29 vector and expressed in Rosetta2pRARE2 cells as done previously for other alphavirus capsid proteins (17, 18, 48, 54, 59). For ease of purification, we included a 6-His tag at the N-terminus prior to the first amino acid. Cells were grown at 37°C, when the OD600 was between 0.4-0.6, 1 mM IPTG was added. Cells were grown for an additional 4 hours. Cells were pelleted and resuspended in 20 mM sodium phosphate buffer containing 2μg/ml leupeptin, 2 μg/ml aprotinin, and 1mM PMSF to a final cell density of 0.06-0.08 g/ml (53). Samples were lysed using a cell cracker or a sonicator, sodium chloride was added to a concentration of 500 mM, and then lysates centrifuged at in a Beckman JA17 rotor at 23000 x *g* or 45 minutes at 4°C. Clarified supernatant was applied to a 5ml HisTrap column (GE Lifesciences) and after washing with 5 to 10 column volumes of 20 mM sodium phosphate, pH 7.4, 500 mM NaCl, and 10 mM imidazole until A280 and A260 returned to baseline, the protein was eluted with a similar buffer but with 800 mM imidazole. For further purification, peak fractions were collected and diluted to 0.25M NaCl with 20 mM sodium phosphate, pH 7.4. Samples were applied to a HiTrap SP column (GE Lifesciences), washed with 20 mM HEPES pH 7.4, 0.25 M NaCl, 5 mM EDTA and then eluted from the column using 20 mM HEPES pH 7.4, 1.3 M NaCl, and 5 mM EDTA. Fractions with an absorbance substantially above background and an 260/280 absorbance ratios less than 0.6 were pooled, concentrated with a centricon 10K concentrator, and buffer exchanged into 20 mM HEPES and 150 mM NaCl. The resulting material served as our stock solution. Protein concentration was determined by measuring absorbance at 280 nm using an extinction coefficient of 39 670 M^-1^ cm^-1^.

### 2.2 High Throughput Screen

High throughput screen assays were performed at the Chemical Genomics Core Facility at IUPUI (Indianapolis, IN). Each compound of the Chembridge 10K library (San Diego, CA; kindly provided by Assembly Biosciences) was prepared as a 1mM stock in 100% DMSO in 384-well plates. We conducted two screens by adjusting the assembly conditions. For the inhibitor screen, we used a final NaCl concentration of 320 mM to achieve approximately 75% assembly of CLPs and focused on compounds that inhibited assembly. In the second screen, we used a final NaCl concentration of 570 mM NaCl to achieve approximately 30% assembly of CLPs and focused on compounds that promoted assembly. The final reaction for each screen contained 0.25 μM of capsid protein, 0.25 μM of oligomer, and 10 μM of compound in a total volume of 25 μl per well in a 384-well plate (Greiner bio one).

For the screens, capsid protein stock was diluted to 0.417 μM in either 534.4 or 951.9 μM NaCl in 20 mM HEPES, pH 7.5 buffer. A 27mer oligomer (5’-TACCCACGCTCTCGCAGTCATAATTCG) was prepared in 20 mM HEPES, pH 7.5 buffer, without any NaCl. The oligomer mixture contained unlabeled 27mer, Cy3-5’ labeled 27mer, and Cy5-5’ labeled 27mer at a 1:2:2 ratio (IDT).

For reactions with compounds, 15 μl of 0.417 μM capsid in the appropriate NaCl solution was first added to each well of a 384-well plates (MultiFlo FA Microplate Dispenser, Bio Tek). Then, 0.25 μl of compound (1 mM stock in DMSO) was added (Freedom EVO 100 liquid handler, Tecan). After a 5-10 minute incubation at room temperature to initiate compound-protein interaction, 10 μl of the 0.625 μM oligomer mixture was added to a final volume of 25 μl. Plates were centrifuged briefly and incubated at room temperature for 2 hours in the dark. Finally, the fluorescence signal was recorded with an EnVision 2102 Multilabel Plate Reader (Perkin Elmer) with monochromators set to 531 nm excitation and 595 nm emission.

Each assay consisted of thirty-two 384 plates and each plate contained a series of controls. One column (16 wells) controlled for assembly in the presence of increasing ionic strength to ensure the fluorescence signal inversely correlated to assembly. A second column had assembly reactions at the salt concentration being used in the assay (320 mM for inhibitors and 570 mM for promoters) to give a baseline fluorescence signal for 75% assembly and 30% assembly, respectively. The third and fourth columns contained reactions with maximal assembly (100 mM NaCl) and minimal assembly (800 mM) to establish the minimum and maximum fluorescence signals, respectively. Also included were capsid protein/no oligonucleotide controls.

### 2.3 High Throughput Data Analysis

HTS data quality was monitored using the standard curve as well as the positive and negative controls on each plate. Data points were normalized by averaging of all the data points in the same position across all plates to eliminate systematic instrumental error on specific wells. We defined potential hits in the following way to minimize random error and allow comparison of chemical structures: hits are 3 sigma (i.e. 3 standard deviations of the signal distribution) away from the average of the primary screen and 2 sigma from the average of the secondary screen.

All analysis was performed using the R project for Statistical Computing (60).

### 2.4 Tissue culture reagents

All tissue culture experiments were performed using the baby hamster kidney cell line BHK-21, here referred to as BHK cells. Cells were passaged in minimum essential medium (MEM) supplemented with non-essential amino acids, penicillin-streptomycin, and L-glutamine in 10% fetal bovine serum (Corning Cellgro, Manassas, VA) and grown at 37°C and 5% CO_2_.

### 2.5 Virus preparation and titering

The CHIK 181/25 virus used in this study was a generous gift from Dr. Terence Dermody’s lab. Nanoluc was cloned into the hypervariable loop of nsP3, after amino acid 490, or after capsid, as described in the literature (61, 62). Infectious virus was generated as described in (63). Briefly, SacI-linearized CHIK-capsid::nLucFM2α cDNA plasmid was transcribed into infectious RNA *in vitro* using a synthetic cap analog and SP6 RNA polymerase (New England BioLabs, Ipswich, MA). RNA was electroporated (1500 V, 25 µF, 200 Ω) into BHK cells resuspended in phosphate-buffered saline (PBS) in a 2-mm cuvette. Upon display of significant cytopathic effect, media were harvested and clarified at 5,000 × *g* for 5 min.

Plaque assays, as described in (63), were used to determine the amount of infectious virus present and was measured as plaque-forming units (PFUs) per mL. Briefly, serial dilutions of the media harvested from virus-infected cells were added to BHK monolayers for 1 h at room temperature while rocking gently. Cells were overlaid with 1% low-melt agarose, 1× complete MEM, and10% fetal bovine serum. At 48 hpi 1 mL of complete MEM with 10% FBS was added to each well. Plaques were detected at 72 hpi by formaldehyde fixation and crystal violet staining.

### 2.6 Compound toxicity in uninfected cells

BHK cells (30,000 cells/well) were plated into a 96-well plate approximately 24 hours prior to adding the compounds. The supernatant was aspirated off and 100μl fresh phenol red-free complete MEM with 10% FBS was added. Then, compounds dissolved in DMSO were added to a final concentration of 10 μM. Plates were incubated for 24 hours at 37°C with 5% CO2. Then, media was aspirated and we assessed ATP levels in cell lysates using the CellTiter-Glo® Luminescent Cell Viability Assay (Promega). Briefly, 30 μL of cell-titer glow reagent was added to each well and, after 10 minutes, the luminescence was measured using a Synergy H1 plate reader (BioTek).

### 2.7 Antiviral assays

To test if compounds inhibited infectious virus production, 300,000 BHK cells/well were plated into 12-well plates approximately 24 hours prior to virus infection. The day of the infection, the supernatant was aspirated and the CHIK capsid:nLucFM2α virus was added at an MOI=0.01 in a final volume of 200 μL. After a 1 hour adsorption period at room temperature, cells were washed twice with PBS to remove unabsorbed virus and then 1 mL of media was added. Compounds were added either during absorption and/or after washing the cells with PBS depending on the experiment. For mock-infected cells, DMSO alone was added. For dose response assays, serial dilutions of each compound were made using DMSO so that the same volume of DMSO was added to culture media. Twenty-four hours after completion of adsorption, supernatants were collected and viral titers determined by plaquing the cells on BHK cells as described above.

### 2.8 qPCR to measure particle numbers

At 24 hrs post infection, supernatants from infected BHK cells were collected and clarified (5000 x g for 5 min at room temperature). Virus concentration was determined by RT-qPCR using 5 μl of each supernatant as template for cDNA synthesis and the method described (64). Primers used were Forward: 5’ggaataaagacggatgatagc and Reverse: 5’ggtcgggaatgaaatttttcc. The absolute quantities of viral RNAs were determined using a standard curve of *in vitro* transcribed RNA.

### 2.9 Nucleocapsid core isolation from infected cells

Cytoplasmic cores from virus-infected cells were isolated as described previously (65). Briefly, four 100 mm dishes of BHK cells were infected with CHIK at a MOI=0.1. Virus was allowed to adsorb for one hour and then cells were washed twice with PBS. To two plates, serum-free media and 1% DMSO was added, and to the other two plates, 1 μM 4BSA (in DMSO) in serum-free media was added. Cells were harvested 24 hours post infection, pellets were washed with ice-cold PBS, resuspended in 1 ml of TNE buffer (10 mM TrisCl, pH 7.5 10 mM NaCl, and 20 mM EDTA). Cells were incubated on ice for 20 minutes. The nuclei were removed by spinning at 1000 x *g* for 10 minutes at 4C. Supernatants were loaded on 10-40% sucrose gradients in TNE+0.1% Triton X-100 buffer. Gradients were spun in SW41 rotor for 2.5 hours at 32,000 x *g* at 15°C. Fractions collected and analyzed by western blot (65).

## 3. Results and Discussion

### 3.1 A high throughput screen to identify compounds that modulate core-like particle assembly

We chose to identify antiviral compounds that target the CHIKV capsid protein because the capsid protein must make several crucial interactions to generate infectious virus. The core protects the viral RNA genome during virus transmission, but must dissociate under the correct physiological conditions for virus replication (46). In addition, the properly assembled core interacts with the cytoplasmic domain of the viral glycoprotein E2 (66) suggesting the assembly of the core is critical for particle budding. *In vitro* assembled CLPs are structurally and functionally similar to nucleocapsid cores from infected cells (50, 51, 54) making them ideal systems for studying core assembly (48, 49, 52, 53, 59, 67–69).

CLPs can be assembled *in vitro* by mixing equimolar ratios of capsid protein and 27mer DNA oligomers in the presence of 100 mM NaCl, as previously demonstrated (52). For those experiments 3 μM capsid protein and 3 μM oligonucleotide were used. CLPs formed rapidly at room temperature and were visualized by dynamic light scattering, gel shift assay, and transmission electron microscopy (48, 53, 54). To examine CLP assembly in a high throughput format we used a fluorescent output: Cy3- and Cy5-labeled 27mer oligomers were used as cargo and DNA packaging was observed by fluorescence resonance energy transfer (FRET) for identification and quantification of CLP assembly (57). When the labeled oligomers are incorporated into core-like particles, the fluorophores are forced into close proximity to one another and FRET occurs, resulting in a decrease in the Cy3 emission signal and a corresponding increase in the Cy5 emission signal (Figure 1). We tested different ratios of unlabeled:Cy-3 labeled:Cy5-labeled 27mer oligomer and found the ratio of 1:2:2, respectively, provided the largest fluorescence range while minimizing self-quenching due to Cy3-Cy3 and Cy5-Cy5 interactions. We observed that the decrease in Cy3 fluorescence had a greater dynamic range than the increase in Cy5 fluorescence, thus we used the change in the Cy3 fluorescence as a read-out for oligo packaging and CLP assembly. We found that 1.5 μM capsid protein and 1.5 μM total oligomer provided a strong signal to discriminate between packaged and unpackaged DNA. This ratio still maintained the 1:1 mole ratio while reducing the concentrations of capsid protein and oligomer used.

**Figure 1.**
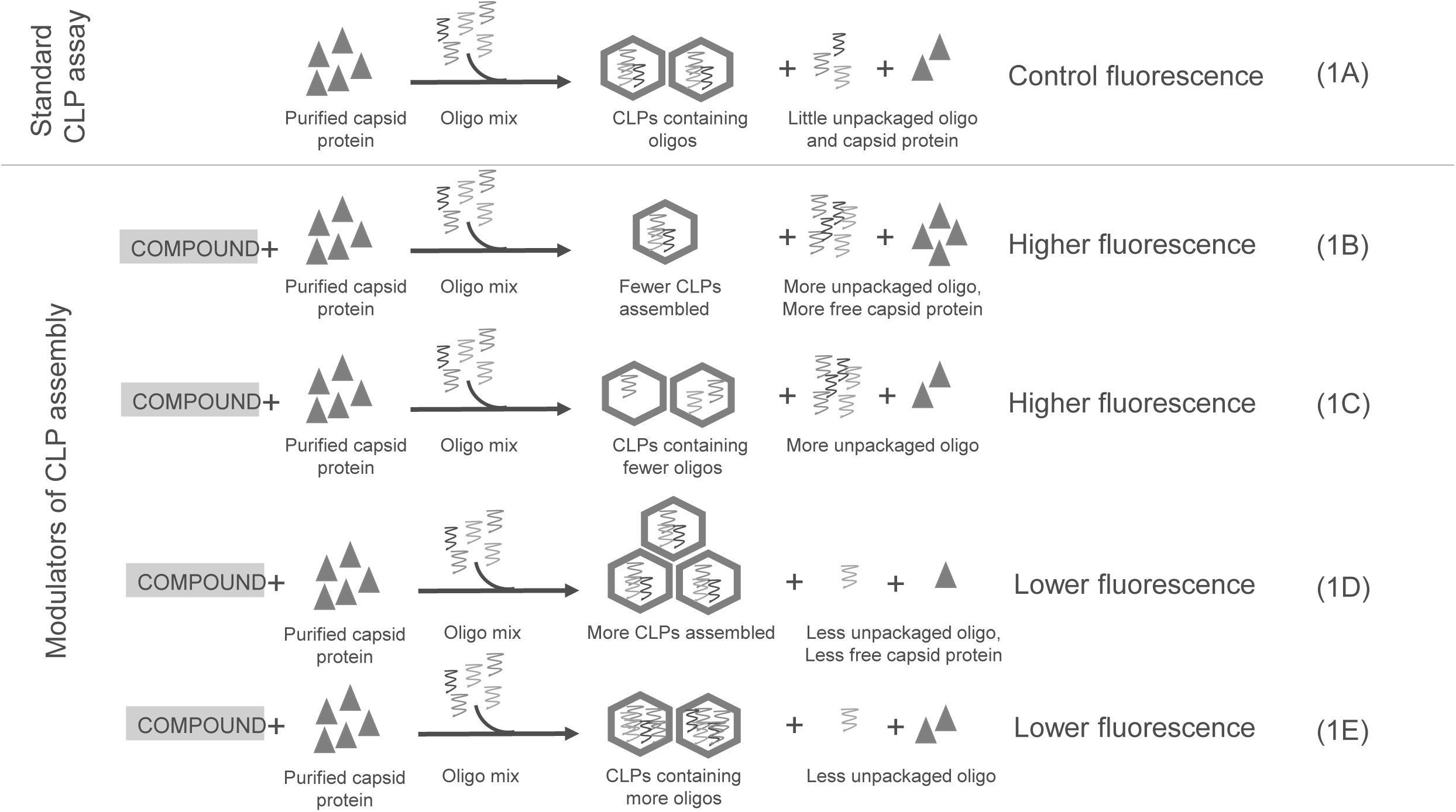
Schematic of the expected outcomes of the FRET-based assay for nucleic acid packaging in CLPs. (A) CLP assay using purified capsid protein and fluorescently labeled DNA oligos that provides the baseline fluorescence. (B-E) Expected outcomes of how fluorescence signal could increase representing less overall oligo packaged (B and C) or decrease representing more overall oligo packaged (D and E) in the presence of added compound.

We used this FRET-based *in vitro* assay (48, 53, 54, 57) to screen 10,000 compounds for their ability to affect fluorescence as a proxy for their ability to modulate CLP assembly. We anticipated the following outcomes (Figure 1). Compared to a control, compounds could inhibit CLP formation resulting in more free nucleotide and higher fluorescence (Figure 1B); they could promote assembly in the absence or reduced amount of DNA also resulting in higher fluorescence (Figure 1C); they could enhance the overall amount of assembly resulting in more oligonucleotide packaging and lower fluorescence (Figure 1D); they could increase the amount of oligos packaged in a CLP, also resulting in lower fluorescence (Figure 1E). Our screen, without other assays, does not determine which scenario is occurring, merely that there is a change in the CLP assembly process.

To maximize our chances of identifying assembly enhancing and inhibiting compounds we screened compounds under two different ionic strength conditions. In the first screen, we used low ionic strength conditions where packaging was robust and fluorescence suppressed.

Compounds that decrease the packaging of oligomers, thereby raising the fluorescence signal from Cy3, would be readily identified (Figures 1B and 1C). In the second screen, we used moderate ionic strength conditions where packaging was reduced and fluorescence higher; this was used to identify compounds that decrease the fluorescence signal from Cy3, suggesting an increase in the packaging of oligomers (Figures 1D and 1E).

To test the core modulation properties of the compounds, CLP assembly was initiated by adding the oligomer mixture to purified capsid protein and 10 μM compound in the appropriate ionic strength buffer. In the lower ionic strength screen, we found 70 molecules out of 10,000 that resulted in an increase in Cy3 fluorescence by at least 3 standard deviations from the mean fluorescence (Figure 2A, dark blue line, +3σ). In the moderate ionic strength screen, we found 36 compounds that decreased Cy3 florescence by at least 3 standard deviations from the mean florescence (Figure 2B, dark green line, -3σ). From our two screens we had at least 106 compounds with CLP modulating properties *in vitro* (Figures 2A and 2B).

**Figure 2.**
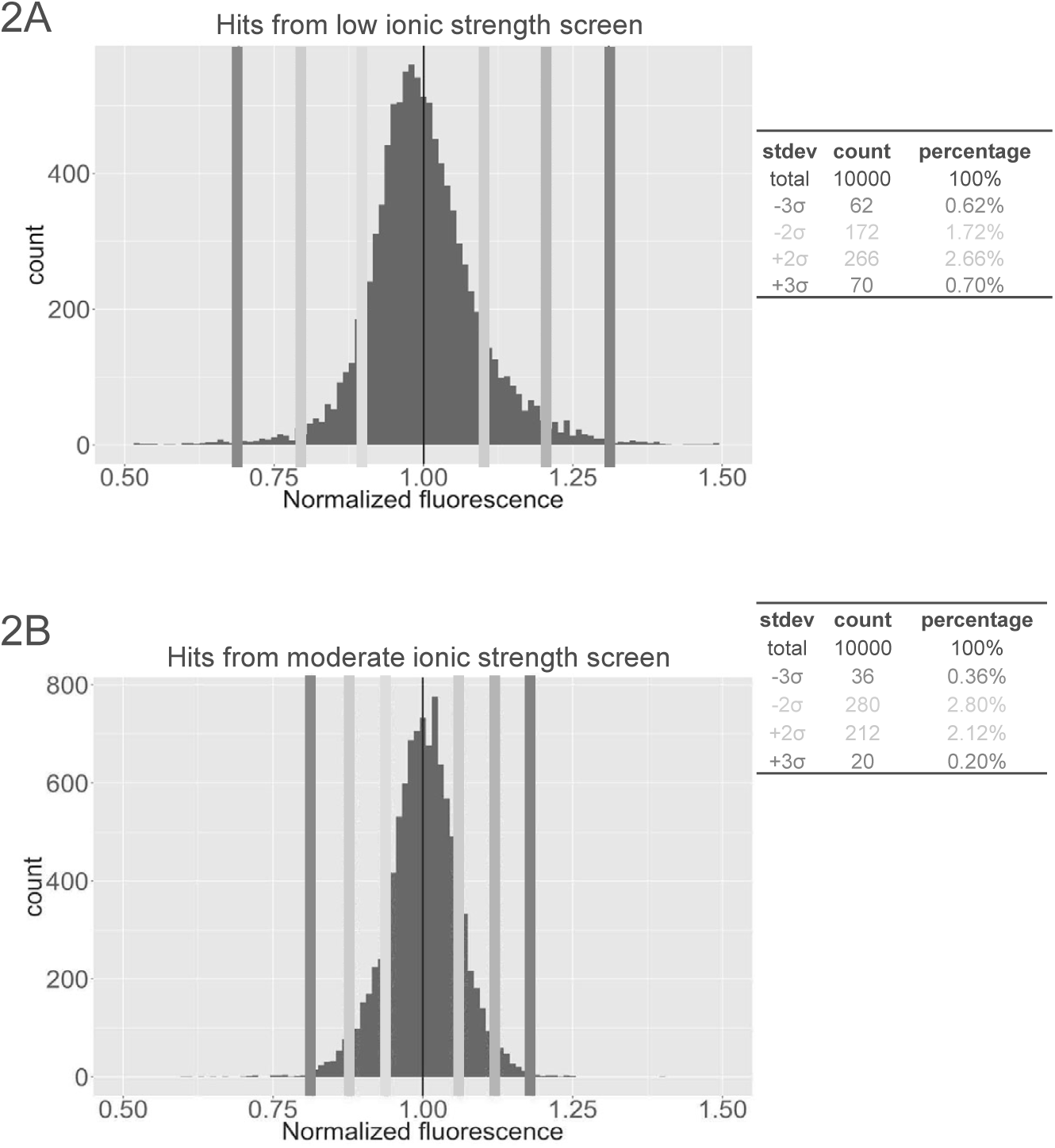
Distribution of normalized fluorescence values from both screens. (A) Screen performed at low ionic strength and (B) at moderate strength. Increased fluorescence values 10, 20, and 30 from the average shown in shades of blue (lightest to darkest, respectively) and decreased fluorescence values with 10, 20, and 30 from the average shown in shades of green (lightest to darkest, respectively). Actual number of hits and percentage of the total shown to the right of each histogram.

There were 19 compounds that increased fluorescence in both the low and moderate ionic strength screens (Figure 3A). Since the moderate ionic strength screen was not as sensitive to detecting increased fluorescence, we looked for compounds that increased fluorescence by 2 standard deviations in this screen that also increased fluorescence by 3 standard deviations in the low ionic strength. In a similar process, 26 compounds that increased fluorescence were identified in both screens (Figure 3B).

**Figure 3.**
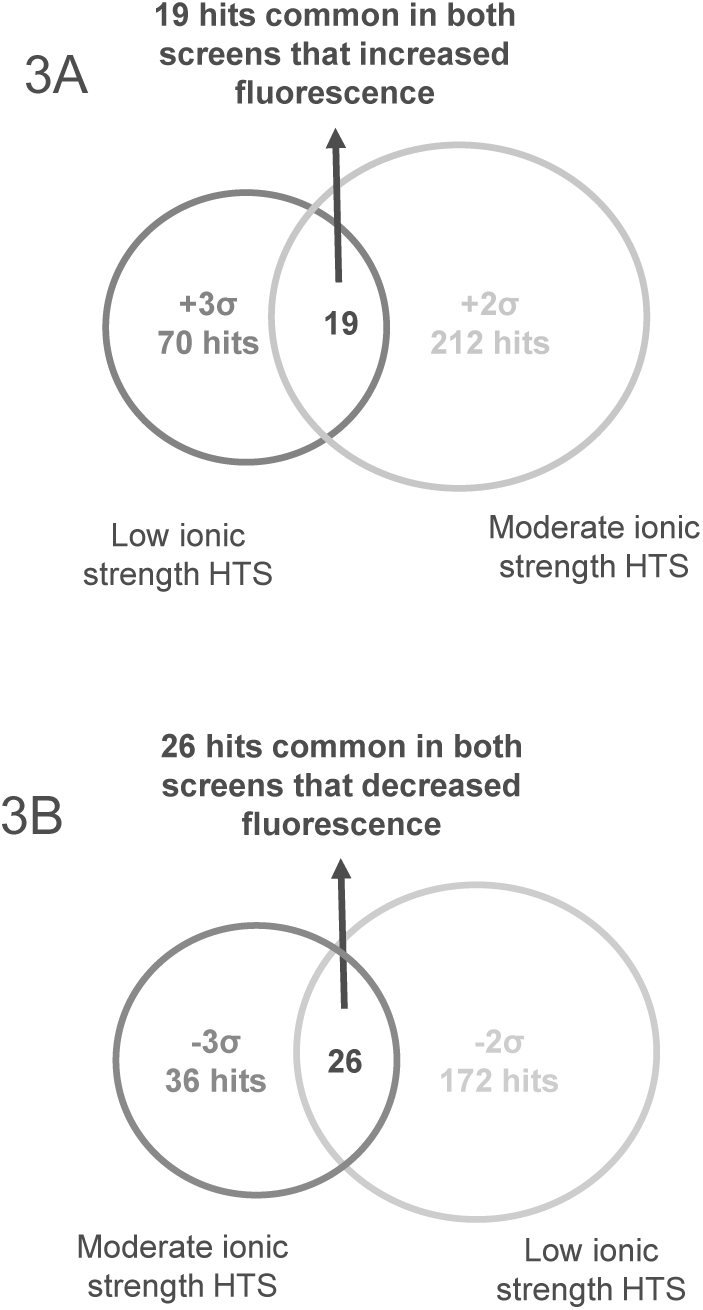
Forty-five modulators of CLP assembly that were selected for further study. (A) 19 compounds identified in both the low and moderate ionic strength screen increased fluorescence during CLP assembly. Compounds 30 above the normalized fluorescence in the low ionic strength screen was compared to compound 20 above the normalize fluorescence in the moderate ionic strength. (B) 26 compounds that decreased fluorescence identified in both screens. Here, the cut-off of 30 was used in the moderate ionic strength and 20 from the low ionic strength screen.

We determined the efficacy of the 45 putative antiviral compounds based on *in vitro* dose response of CLP assembly (Supplemental Figure 1). These assays were performed using standard conditions of 1.5 μM capsid protein, 1.5 μM oligomer, and either 320 mM or 570 mM NaCl. The AC50 (defined as half-maximal activity concentration) of the compounds ranged from 1 to 9 μM. Based on efficacy, 12 compounds, six that increased fluorescence and six that decreased fluorescence, were selected for further characterization.

### 3.2 Production of CHIKV infectious particles is reduced with core modulating compounds

The screen to identify CLP assembly modulators was performed using purified components to maximize the likelihood of finding a compound that interfered with capsid protein-oligo or capsid protein-capsid protein interactions rather than an off-target cellular effect. Now we wanted to determine if the compounds identified as core modulators *in vitro* were also effective during viral infections in cell culture.

The 12 selected compounds were tested for toxicity and antiviral efficacy. To test toxicity, we cultured BHK cells in the presence of 10 μM of each compound for 24 hours and then (i) assessed cell health by measuring cellular ATP levels using a commercial luciferase-based assay and (ii) assessed cellular morphology by microscopy. The 24-hour time point was selected because CHIKV-infected BHK cells show cytopathic effect at this time post-infection. At 10 μM, eight compounds were tolerated well by cells and showed ATP levels similar to control cells treated with DMSO (Figure 4). In contrast, compounds MA, NTU, NDBA, and 4PBSA were detrimental to cells based on reduced levels of ATP and altered cellular morphology.

**Figure 4.**
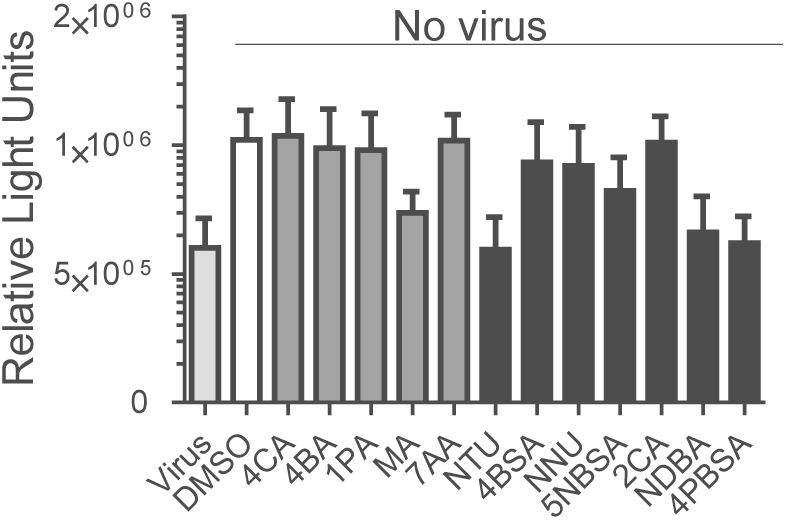
Toxicity assessment of compounds in cell culture. BHK cells were incubated with 10 μM of compound only for 24 hours and then ATP levels were assessed using Cell Titer Glow. Compounds that increase fluorescence, possible assembly inhibitors, are shown with gray bars; compounds that decrease fluorescence, possible assembly enhancers, are shown with black bars. Cells incubated with virus and DMSO served as positive and negative controls, respectively. These data are the averages of 3 independent experiments performed in triplicate. Error bars indicate the standard deviation

Based on these findings, we only tested the eight non-toxic compounds (4CA, 4BA, 1PA, 7AA, 4BSA, NNU, 2P, and 2CA) for their ability to inhibit infectious CHIK virus production. We performed a series of experiments where the time at which the compounds were added was varied relative to CHIK virus adsorption: BHK cells were incubated with the compounds continuously (Figure 5A), only after viral absorption (Figure 5B), or only during viral absorption (Figure 5C). In all cases, supernatants from each of these experiments were collected 24 hours post virus infection and assessed for infectious virus by plaque assays. All eight of the compounds had anti-viral activity (Figure 5), albeit to differing extents depending on when the compound was added during the infection.

**Figure 5.**
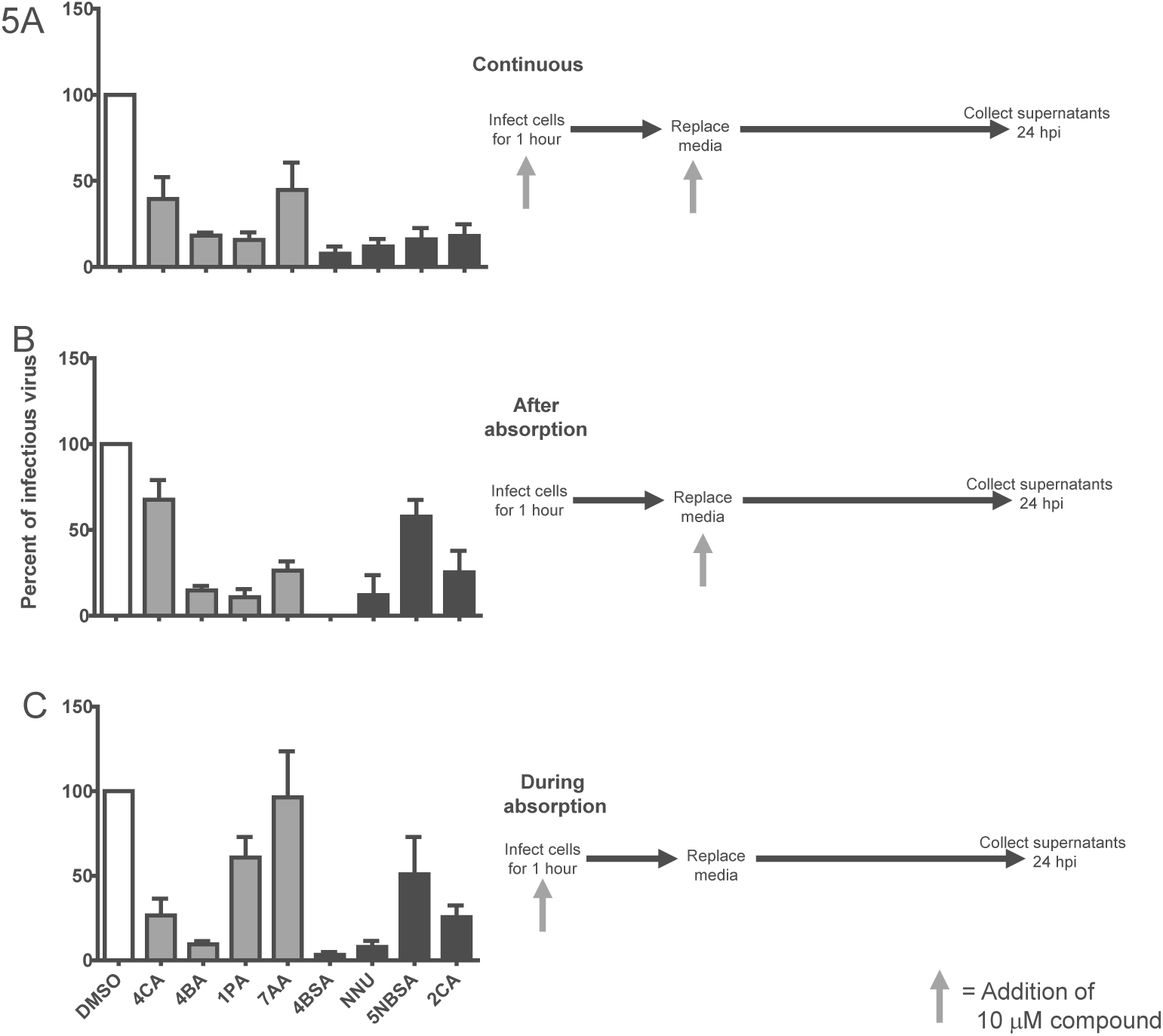
Antiviral activity of compounds as a function of their time-of-addition. BHK cells were infected with CHIKV and 10 μM of compound was added (A) continuously, (B) after virus absorption, or (C) during absorption of the virus. In left panel, compounds that increase fluorescence are shown with gray bars; compounds that decrease fluorescence are shown with black bars. In the experimental flowchart (right panels), the gray arrow indicates when compound was added. Supernatants were collected 24 hpi and plaque assays were used to determine the amounts of infectious virus. These data are averaged from three independent experiments. Error bars indicate the standard deviation.

When 10 μM of compound was incubated with CHIKV-infected BHK cells continuously (Figure 5A), all eight compounds inhibited infectious virus production by at least 50% when compared to the DMSO-treated control cells. Compound 4BSA was the most potent inhibitor of infectious virus production and suppressed viral titer by at least 90%. When compounds were added to cells after virus absorption (Figure 5B), they all reduced the amount of infectious virus produced by at least 50%, except for 4CA and 5NSBA. In the final time-of-addition experiment, we treated BHK cells with compound for only 1 hour during absorption of the virus (Figure 5C) and found that compounds 5NSBA, 1PA, and 7AA were much less effective and no longer inhibited production of infectious virus by 50%.

The eight compounds that exhibited antiviral activity were identified from the screen that only contained capsid protein and oligomer. Therefore, we hypothesized the compounds primarily interfere with an aspect of core assembly or disassembly. We cannot say if the antivirals interfere with capsid-nucleic acid, capsid protein-capsid protein, or both types of interactions. The time of addition experiments may suggest whether the compounds could affect early steps in infection (e.g. disassembly) or late steps (e. g. assembly) or at multiple steps. We hypothesize that compounds that led to higher fluorescence inhibit new core assembly and thus limit production of infectious virus. Compounds 1PA and 7AA were consistent with this idea.

They were only an effective antiviral if added continuously or after virus absorption (Figure 5A, 5B), and much of the antiviral activity was lost if this compound was added only during virus absorption (Figure 5C). Compounds 4BA and 4CA were antivirals regardless of time of addition. Compounds that led to lower fluorescence were hypothesized to inhibit nucleocapsid core disassembly, an earlier part of the virus life cycle. Our results show 4BSA, NNU, and 2CA were effective antivirals even when added only during virus absorption (Figure 5C) consistent with these compounds acting during an early step of the viral life cycle. 5NBSA worked the best when added continuously, perhaps working post-disassembly.

Plaque assays reproducibly showed that eight of the compounds identified in our screen inhibited production of infectious virus (multiple independent virus preparations) (Figure 5). Many high throughput screens that identify new antiviral compounds use reporter-based assays to quickly look for a 50-75% reduction in reporter activity (8–11). We initially sought to confirm the antiviral activity of all our compounds by using luciferase reporter viruses. We inserted luciferase into either a hypervariable loop of nsP3 (nsP3:luciferase) or after capsid protein (capsid:luciferase) (61, 62). CHIKV containing the luciferase reporter was added at an MOI=0.01 and rocked for 1 hour. Then the cells were washed and each of the eight compounds was added with fresh media. After 6 or 9 hours when using the nsP3:luciferase or capsid:luciferase virus (when maximum signal was observed), cells were lysed and luciferase activity determined. In our hands, most of the eight antiviral candidates failed to yield consistent luciferase activity results, thereby making it difficult to estimate the effect of the compounds (Supplemental Figure 2). Only 4BSA consistently and reproducibly inhibited the luciferase activity while the other antiviral compounds found in our screen had much wider variation. We found the reporter assay was biased towards identifying compounds that would inhibit entry and replication since time points were taken relatively early in the viral lifecycle. These data suggest that compounds that target the later stages of assembly such as core assembly, particle assembly, and have subsequent effects in particle spread, may be missed in this type of reporter screen.

### 3.3 Efficacy of antiviral compounds in CHIKV infection

We next tested the dose-dependence inhibition of our most inhibitory compounds (4BA, 1PA, 4BSA, and NUU) to estimate their EC50 or the half-maximal effective concentration. Following the addition scheme of Figure 5B, cells were treated with different concentrations of compound after virus adsorption for one hour. EC50 values for 4BA was 0.94 μM, 1PA was 1.19 μM, 4BSA was 0.73 μM, and NUU was 1.23 μM (Figure 6). When we compare these values to what was observed in our initial *in vitro* assays, we see that similar concentrations of compound are required to obtain inhibition of infectious particles in cell culture and CLP assembly *in vitro*. We see 0.94 vs 1.73 μM for compound 4BA, 1.19 vs. 1.75 μM for 1PA, 0.73 vs. 2.44 μM for 4BSA, and 1.23 vs 5.34 μM for NNU when comparing the *in vivo* to the *in vitro* dose responses (Figure 6 and Supplementary Figure 1). The largest difference between *in vivo* and *in vitro* studies was an approximately 4-fold difference for NNU.

**Figure 6.**
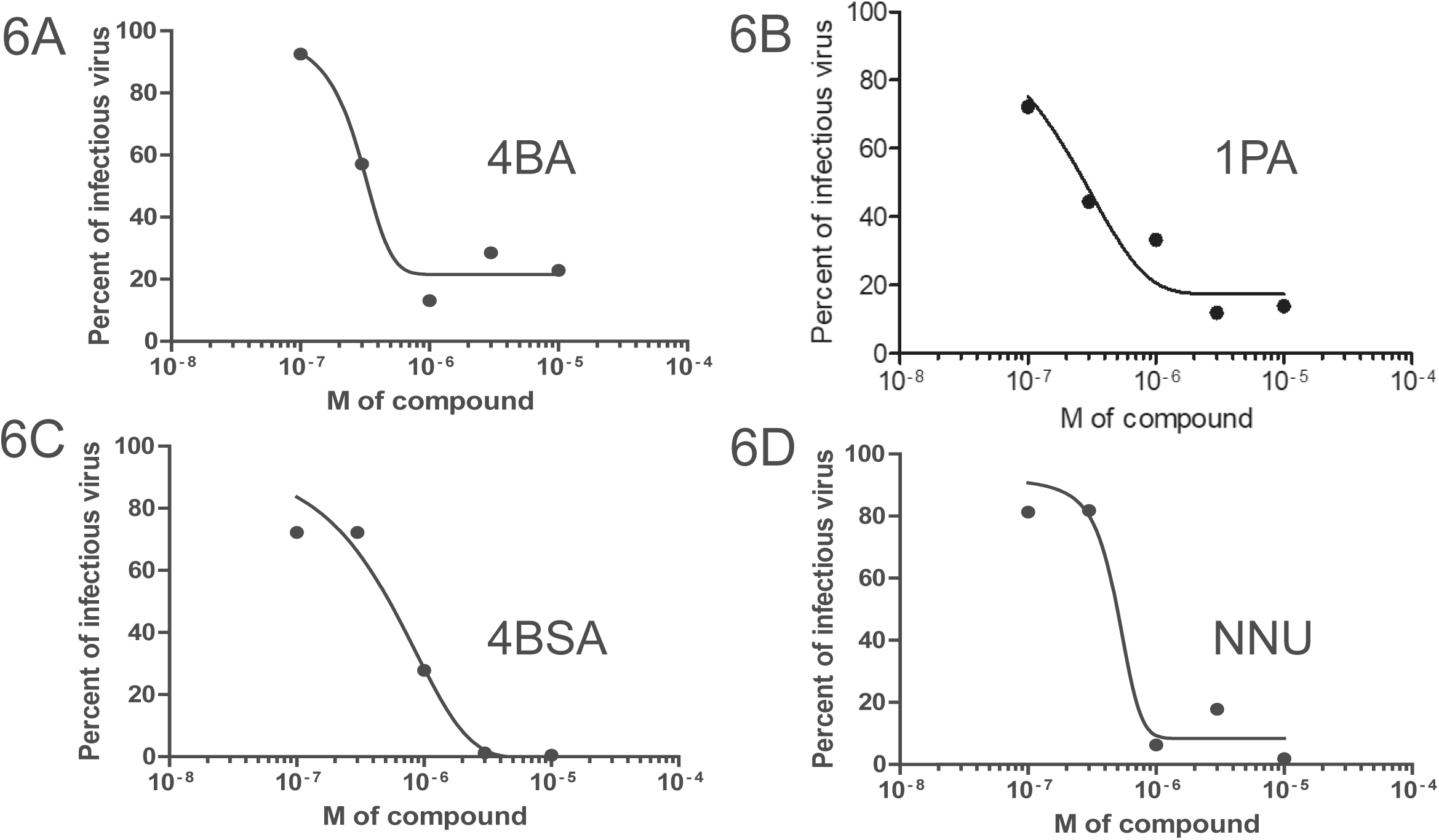
Four compounds have EC_50_ values in the μM range i*n vivo*. BHK cells were infected with CHIKV and 10 μM of compound (A) 4BA, (B) 1PA, (C) 4BSA, and (D) NNU. was added immediately after virus absorption. Supernatants were collected 24 hpi for plaque assays. EC50 values were 0.94 μM for 4BA, 1.19 μM for 1PA, 0.73 μM for 4BSA, and 1.23 μM for NNU. These representative data are from one of three independent experiments. Why do 4BA and 1PA only go to about 20%?

The EC50 observed by our study (Figure 6) indicate that four compounds are promising leads. Several groups have studied the antiviral properties of dioxane and its derivatives which bind in the hydrophobic pocket of capsid protein and subsequently interfere with the capsid-E2 interaction (17–19). Like Sharma *et al*. saw with CHIKV and picolic, we observed the best inhibition when 4BSA and virus were added simultaneously. However, they reported that 2 mM of picolic acid was needed to inhibit infectious virus production by 50-60% (19); in contrast, <2μM of the four compounds identified by our study was needed to reduce infectious virus production by 50%.

### 3.4 Core assembly is targeted by compound 4BSA

The four compounds that we focused on reduced the number of infectious particles released during a virus infection. Their effectiveness depends on the time of their addition, and their EC50 values range from 0.7-1.25 μM. Our initial screen looked at modulators of core assembly. Our best inhibitor, based on dose response and inhibition of infectious virus, was 4BSA. To determine if 4BSA inhibited core assembly in CHIKV-infected cells, we examined lysate from infected cells with and without compound via centrifugation through a sucrose gradient (65).

During a CHIKV infection, nucleocapsid cores form in the cytoplasm and are then enveloped by the viral glycoproteins and bud from the cell. Fractions from the gradient were analyzed by western blot probing for capsid proteins. Virus-infected cells show capsid protein at the top and midway through the sucrose gradient (Figure 7, left panel), consistent with free protein and assembled cytoplasmic cores (65). In contrast, cells treated with 4BSA did not form detectable amounts of cores (Figure 7, right panel). The low overall abundance of capsid protein in cells treated with virus and 4BSA could be the cumulative result of 4BSA activity interfering with the initial disassembly of the virus core, reducing viral replication and new virion propagation, and decreasing amounts of viral spread.

**Figure 7.**
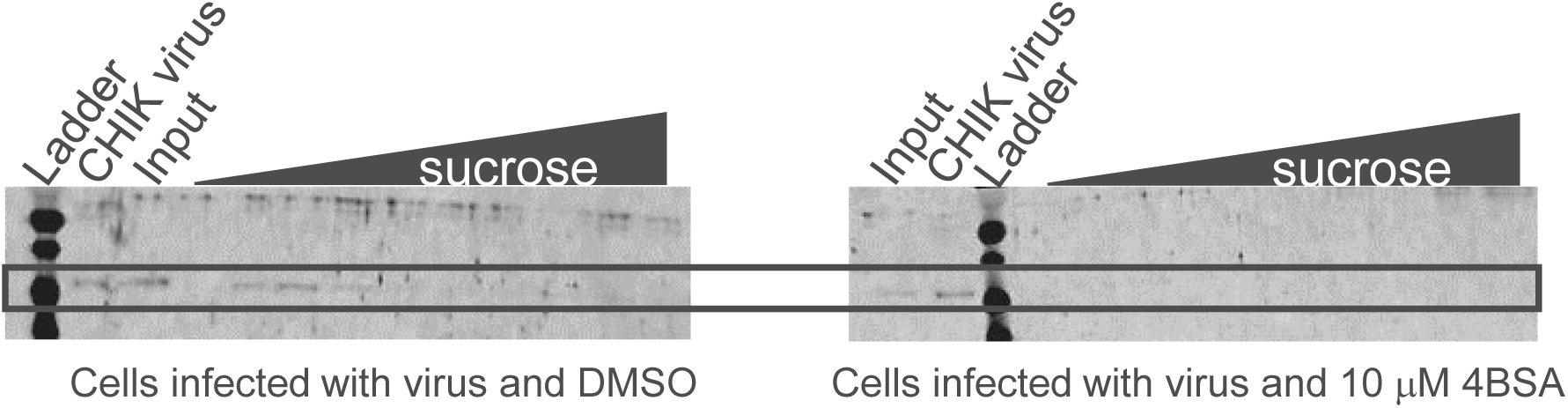
4BSA inhibits nucleocapsid core assembly in CHIKV-infected cells. BHK cells were infected with CHIKV and either DMSO (left) or 10 μM of 4BSA (right) was added immediately after virus absorption. Cells were lysed and nucleocapsid cores were isolated by density centrifugation. Fractions were collected and assessed for the presence of capsid protein by western blot. These representative data are from one of two independent experiments.

**Figure 8.**
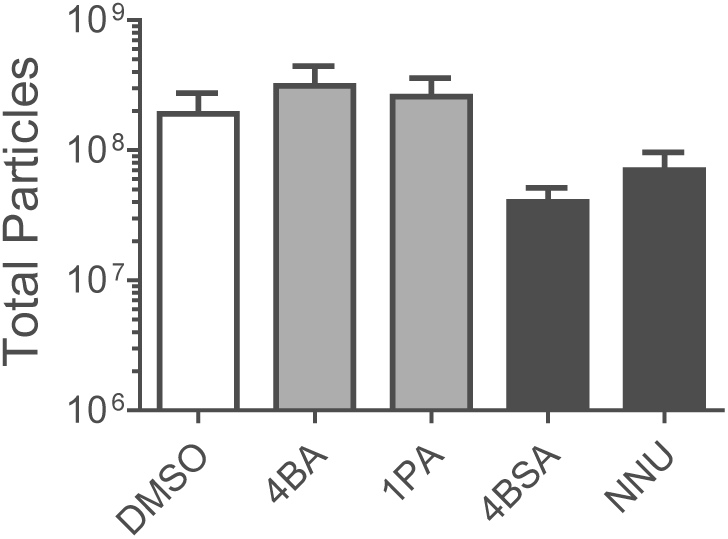
Total number of virus particles released are slightly affected by the presence of compound. BHK cells were infected with CHIKV and 10 μM of compound was added immediately after virus absorption (similar to Figure 5B). Compounds that increase fluorescence are shown with gray bars; compounds that decrease fluorescence are shown with black bars. Supernatants were collected 24 hpi for qPCR studies. These data are averaged from three independent experiments. Error bars indicate the standard deviation.

Viral genomes are present in both infectious and non-infectious particles. To determine if the total amount of particles released was altered in the presence of these compound, we determined the genome copies in released CHIKV. CHIKV was added to cells and after adsorption, compound was added, similar to Figure 5B. After 24 hours, media was collected and assayed by qRT-PCR (Figure 6). All four compounds affected the yield of total particles by less than one-log though infectious particles decreased by more than 1-log. These results suggest that viral genome are packaged in the presence of compound but the genome is not able to infect new cells.

## 4. Conclusions

In summary, we used a novel FRET-based assay to screen 10,000 compounds for their ability to inhibit or promote assembly of core-like particles. Of the initial 106 hits, eight compounds were further assessed due to their potent *in vitro* dose response and their lack of cellular toxicity. From this group, we identified four compounds that could reduce production of infectious virus by at least 1-log and at least one of these compounds interferes with nucleocapsid core assembly *in vivo*. These results indicate that compounds with the ability to modulate core-like particle assembly also have potent antiviral activity. Our work adds to the growing body of evidence that suggest that targeting the capsid protein is an effective strategy to find new antivirals.

## 5. Acknowledgments

We thank members of the Mukhopadhyay and Zlotnick lab for scientific discussions. This work was supported by Assembly Biosciences (SM). AZ associated with biotechnology companies that focus on Hepatitis B virus.

## Supplemental figures

**Supplemental Figure 1.**
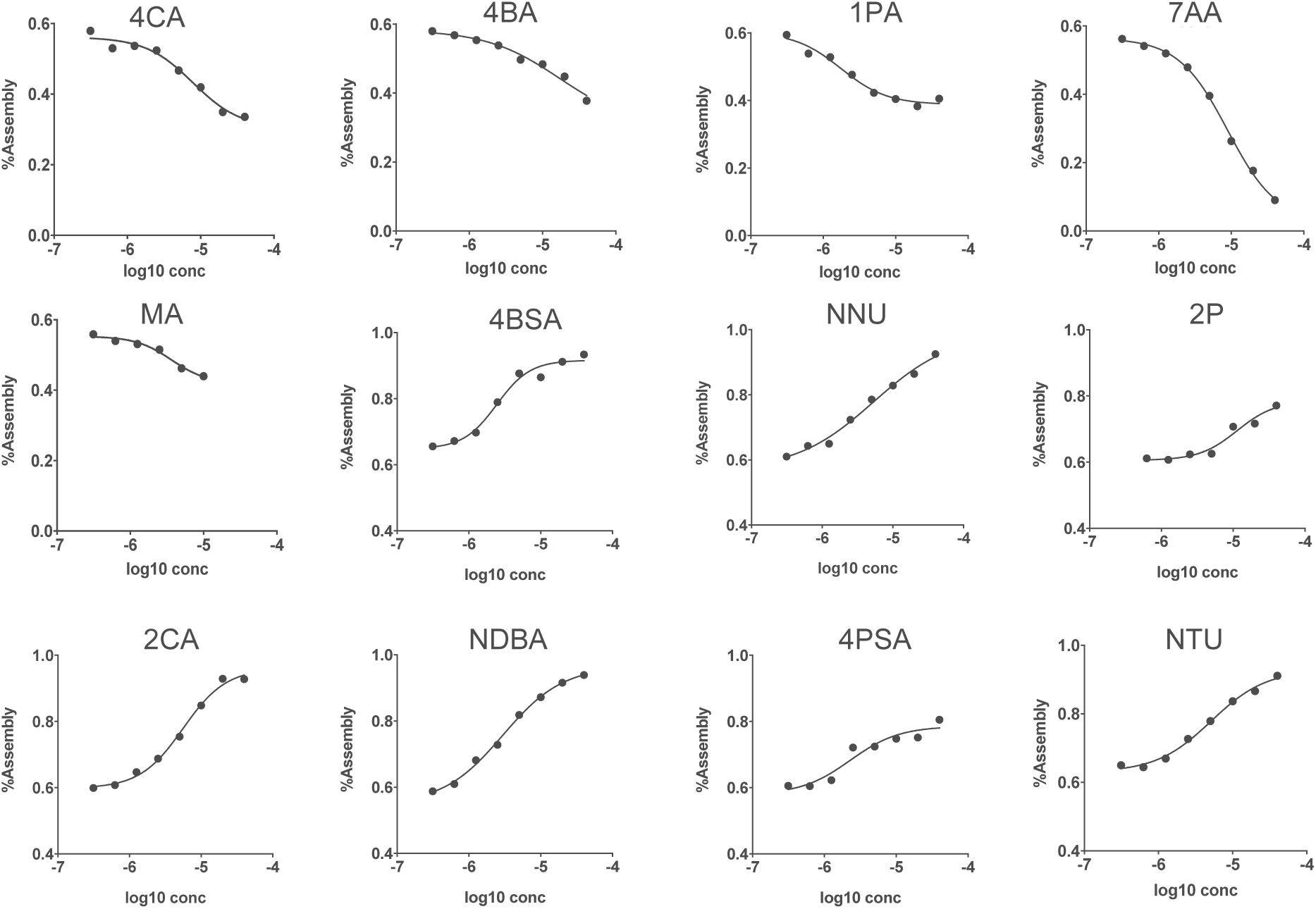
Forty-five of the CLP modulators identified in the HTS were validated by determining CLP assembly *in vitro* as a function of compound concentration. An increase in fluorescence signal correlated to low assembly and a decrease in fluorescence signal correlates to high assembly. Dose response of compounds that increase fluorescence were tested at 320 mM NaCl and dose response of compounds that decrease were tested at 570 mM. The twelve compounds shown here were selected to be further tested in tissue culture assays using intact CHIK virus.

**Supplemental Figure 2.**
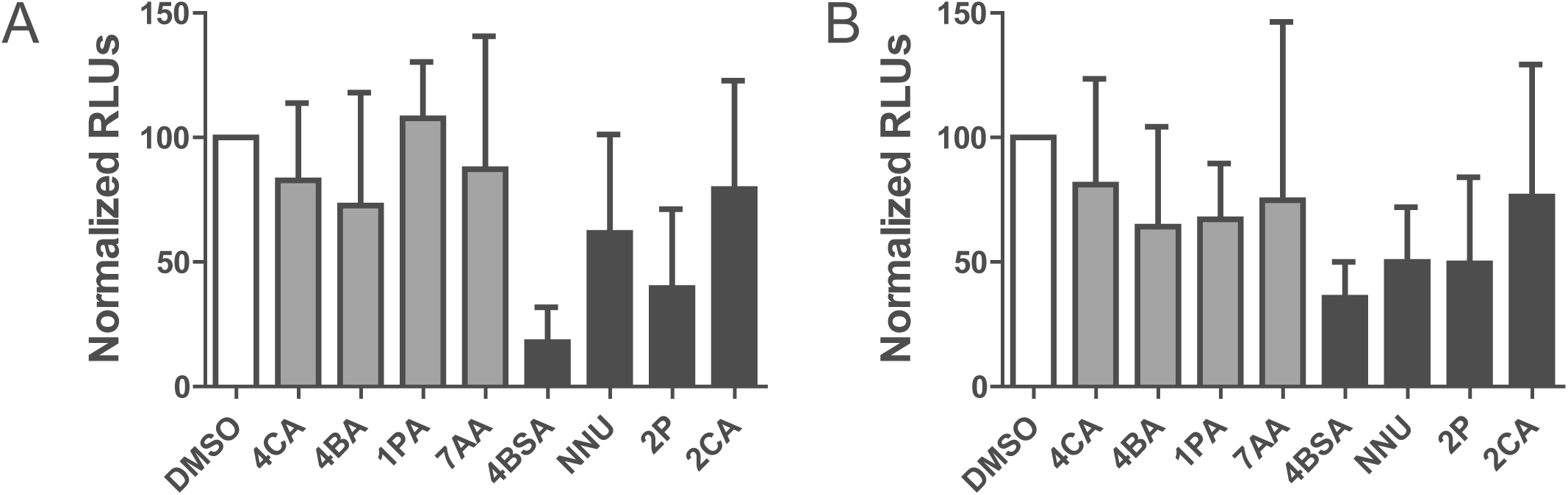
Sensitivity of reporter gene assays to compounds identified by the assembly assay. BHK cells were infected with CHIKV nsP3:luciferase (A) or capsid:luciferase (B) and 10 μM of compound was added immediately after virus absorption. Cells lysates were assessed for luciferase either 6 hpi (A) or 9 hpi (B). These data are averaged from 3 independent experiments; each performed in triplicate. Medium gray and black bars represent compounds identified in the inhibitor or promoter screen, respectively.

